# Regulation of ribosomal protein synthesis in mycobacteria: autogenous control of *rpsO*

**DOI:** 10.1101/2021.02.16.431553

**Authors:** Leonid V Aseev, Ludmila S Koledinskaya, Oksana S Bychenko, Irina V Boni

## Abstract

Autogenous regulation of ribosomal protein (r-protein) synthesis plays a key role in maintaining the stoichiometry of ribosomal components in bacteria. Our main goal was to develop techniques for investigating the r-protein synthesis regulation in mycobacteria, Gram-positive organisms with a high GC-content, which has never been addressed. We started with the *rpsO* gene known to be autoregulated by its product, r-protein S15, in a broad range of bacterial species. To study the in vivo regulation of *rpsO* from *Mycobacterium smegmatis* (*Msm*), we first applied an approach based on chromosomally integrated *Msm rpsO’*-’*lacZ* reporters by using *E. coli* as a surrogate host. The β-galactosidase assay has shown that mycobacterial *rpsO* expression is feedback regulated at the translation level in the presence of *Msm* S15 in trans, like in *E. coli*. Next, to overcome difficulties caused by the inefficiency of mycobacterial gene expression in *E. coli*, we created a fluorescent reporter system based on *M. smegmatis*. To this end, the integrative shuttle plasmid pMV306 was modified to provide insertion of the *Msm* or *Mtb* (*M. tuberculosis*) *rpsO-egfp* reporters into the *Msm* chromosome, and a novel *E. coli*-mycobacteria replicative shuttle vector, pAMYC, a derivative of pACYC184, was built. Analysis of the eGFP expression in the presence of the pAMYC derivative expressing *Msm rpsO vs* an empty vector confirms the autogenous regulation of the *rpsO* gene in mycobacteria. Additionally, we have revealed that the mycobacterial *rpsO* core promoters are rather weak and require upstream activating elements to enhance their strength.

**IMPORTANCE:** Bacterial ribosomes are targets for a majority of as-yet reported antibiotics, hence ribosome biogenesis and its regulation are central for development of new antimicrobials. One of the key mechanisms regulating ribosome biogenesis in bacteria is the autogenous control of r-protein synthesis, which has been so far explored for *E. coli* and *Bacillus spp*. but not yet for mycobacteria. Here, we describe experimental approaches for in vivo analysis of mechanisms regulating r-protein synthesis in mycobacteria, including *M. tuberculosis*, and show, for the first time, that the autogenous control at the translation level is really functioning in these microorganisms. The developed system paves the way for studying various regulatory circuits involving proteins or sRNAs as mRNA- targeting *trans*-regulators in mycobacteria as well as in other actinobacterial species.

## INTRODUCTION

Biogenesis of ribosomes in bacteria is energetically costly as it requires the balanced synthesis of three rRNA molecules and multiple ribosomal proteins (r-proteins) in stoichiometric amounts. Among the mechanisms maintaining coordinated synthesis of ribosomal components, the role of autogenous control of r-protein operons is widely recognized (1–4). An ability of r-proteins synthesized in excess over rRNA to inhibit expression of their own mRNAs has been already shown for most *E. coli* r-protein operons (3, 5–8). However, our knowledge of the r-protein - mediated regulation is based mainly on investigations conducted on *E. coli* and its close relatives in γ-proteobacteria or, to a lesser extent, on *Bacilli*, low-GC Gram-positive organisms. Almost no information is available for other bacterial phyla, including Actinobacteria which are Gram-positive organisms with a high GC- content. This phylum comprises many human pathogens, e.g., *Mycobacterium tuberculosis* and *Mycobacterium leprae* in the genus *Mycobacterium*, which highlights the importance of studies on the mechanisms of mycobacterial gene expression and its regulation.

One of the most thoroughly studied case in autogenous regulation of bacterial r-proteins is the *rpsO* gene encoding r-protein S15, a primary r-protein in the assembly of the 30S ribosomal subunit. The details of the S15-mediated autogenous control have been examined in numerous works dedicated to the *rpsO* expression regulation in *E. coli* (9–13) *Bacillus stearothermophilus* (14, 15*), Geobacillus kaustophilus* (16), *Thermus thermophilus* (17), *Rhizobium radiobacter* (18). The *rpsO* regulation in all these cases operates at the translation initiation level through binding of S15 to specific regulatory structures in the 5’ untranslated region (5’UTR) of the *rpsO* mRNA, leading to inhibition of translation either by the ribosome “entrapment” in a non-productive complex (*E. coli*, see 9, 13) or by a direct competition with the ribosome binding (*Th. thermophilus*, see 17). Interestingly, despite a high conservation of S15 across eubacteria, the regulatory RNA structures (translational operators) widely vary both at the primary and secondary structure levels, suggesting a high S15-RNA interaction plasticity (18, 19).

As a rule, operator structures controlling r-protein synthesis in *E. coli* are highly conserved across several families in γ-proteobacteria, and in a few cases more widely (3). In *Bacilli*, up to now r-proteins S15, L20, S4 and L10(L12)_4_ have been experimentally validated as autogenous regulators, and comparative genomic studies revealed significant conservation of the corresponding operator structures in this class of Firmicutes (20, 21). Thus, the conservation of secondary/tertiary structures within 5’UTRs of r-protein mRNAs in as-yet unexplored species may be indicative of the autogenous control. Recently, computational analysis of the *rpsO* 5’UTRs predicted the presence of the conserved structural elements in Actinobacteria, indicating a high probability for the *rpsO* autogenous regulation in this phylum, but this hasn’t yet been confirmed experimentally (18).

To study mechanisms for the control of r-protein synthesis in bacterial species, it is necessary first to design technical approaches allowing the reliable identification of the feedback loops functioning in vivo. Such methods have been developed for *E. coli*, and as a result, a majority of *E. coli* r-protein operons has been already studied (see above). At the same time, for organisms with a high GC-content like Actinobacteria, the adequate in vivo approaches have not yet been assayed. Here, taken the *rpsO* gene as a classic example, we address the regulation of r-protein synthesis in *M. smegmatis* (*Msm*) and *M. tuberculosis* (*Mtb*). We used a previously developed strategy based on chromosomally integrated reporter genes under the control of the *rpsO* regulatory regions and ectopic expression of *Msm* S15 to measure its effect on the reporter expression. This approach allows a quantitative evaluation of the impact of the excess r-protein on efficiency of its own mRNA regulatory region.

First, we used *E. coli* as a surrogate host. An inhibiting effect of *Msm* S15 *in trans* on the *Msm rpsO’-’lacZ* expression provided evidence that regulation of the *Msm rpsO* gene bears close resemblance to that found for *E. coli rpsO*. However, expression of the mycobacterial *rpsO* gene in *E. coli* turned out to be ineffective, indicating that to continue research into other mycobacterial r-protein operons, it is necessary to develop a cognate system based on *M. smegmatis* as an efficient host for mycobacterial gene expression (22). We developed the fluorescence reporter assay by modifying the integrative shuttle vector pMV306 (23) to transfer the reporter constructs *Msm rpsO-egfp* onto the *Msm* chromosome, and to provide expression of *Msm* S15 *in trans*, we built a new *E. coli*-mycobacteria replicative shuttle vector, pAMYC, on the basis of pACYC184. The results of fluorescence measurements demonstrate that the *Msm rpsO* gene is feedback regulated at the translation level, thereby strengthening the data obtained with *E. coli* as a surrogate host. In addition, by using the *Msm* expression system, we demonstrated the *rpsO* autogenous control for *M. tuberculosis*, thus providing evidence that the proposed system works with different mycobacterial species. The developed approach is applicable to investigating the mycobacterial gene expression regulated by the mRNA-targeting *trans*-factors such as RNA-binding proteins or sRNAs.

## RESULTS and DISCUSSION

### A strategy for using *Escherichia coli* as a surrogate host for studying autogenous regulation of mycobacterial r-proteins

Post-transcriptional control of gene expression in Actinobacteria, including protein- or sRNA-mediated riboregulation, is poorly investigated. Our main goal was to develop techniques allowing examination of the r-protein autogenous control in mycobacteria, which has never been addressed. We started with the *rpsO* gene which was shown to be negatively regulated by its product, r-protein S15, in a range of different bacterial species (see Introduction). To study the in vivo regulation of *rpsO* from *M. smegmatis* (*Msm*) as a model mycobacterium, we first applied an approach based on chromosomally integrated *rpsO’*-’*lacZ* reporters by using *E. coli* as a surrogate host. This methodology includes the creation of the *rpsO’-’lacZ* reporter under the control of the *Msm rpsO* regulatory region on the plasmid pEMBLΔ46 (24) and its subsequent transferring onto the chromosome of a specialized *E. coli* strain ENS0 (24) by homologous recombination to provide stable expression from a single-copy reporter gene. The use of *E. coli* as a surrogate host has been previously exploited to study the autogenous control of different r-protein operons from γ-proteobacteria (25, 26, 6, 8) but its application to other bacterial phyla, especially to those with a high GC-content, has not yet been corroborated. Given that transcription and translation machineries of *E. coli* and mycobacteria have both common and significantly divergent features (27–34), it was difficult to predict in advance whether expression of a certain mycobacterial gene in *E. coli* would be effective, like described in (30), or not. This needed to be experimentally verified.

### Comparison of regulatory regions for mycobacterial and *E. coli rpsO* genes

The promoter and translation initiation regions (TIRs) of both the *M. smegmatis* (*Msm*) and *M. tuberculosis (Mtb) rpsO* genes resemble those of *E. coli*, though some details appear to be quite different (Fig. 1 A, B). While the promoter element −10 (consensus TANNNT, see 35) is present in mycobacterial *rpsO*, the consensus region −35 is not readily defined, which is typical of mycobacterial promoters (34, 35). At the same time, both *Msm* and *Mtb rpsO* promoters belong to the class of the extended −10 promoters (TGnTANNNT) which are recognized by *E. coli* RNA polymerase (27, 36), albeit the absence of the conserved TTG in a region −35 (*Msm*) may have a negative impact on the promoter activity (36). The initiator codon is a GUG both in *Msm* and *Mtb rpsO*, while in *E. coli* the *rpsO* coding region starts with an AUG. It is known that a GUG is used more often in mycobacteria than in *E. coli* (34), but given that several *E. coli* genes (e.g., *rpsM* encoding the r-protein S13) show a high expression level with a GUG start codon, a combination of the *rpsO* GUG with a canonic Shine-Dalgarno (SD) element (GGAG in *Msm, Mtb* and *E. coli*) may be estimated as recognizable by *E. coli* ribosome during translation initiation.

**Figure 1.**
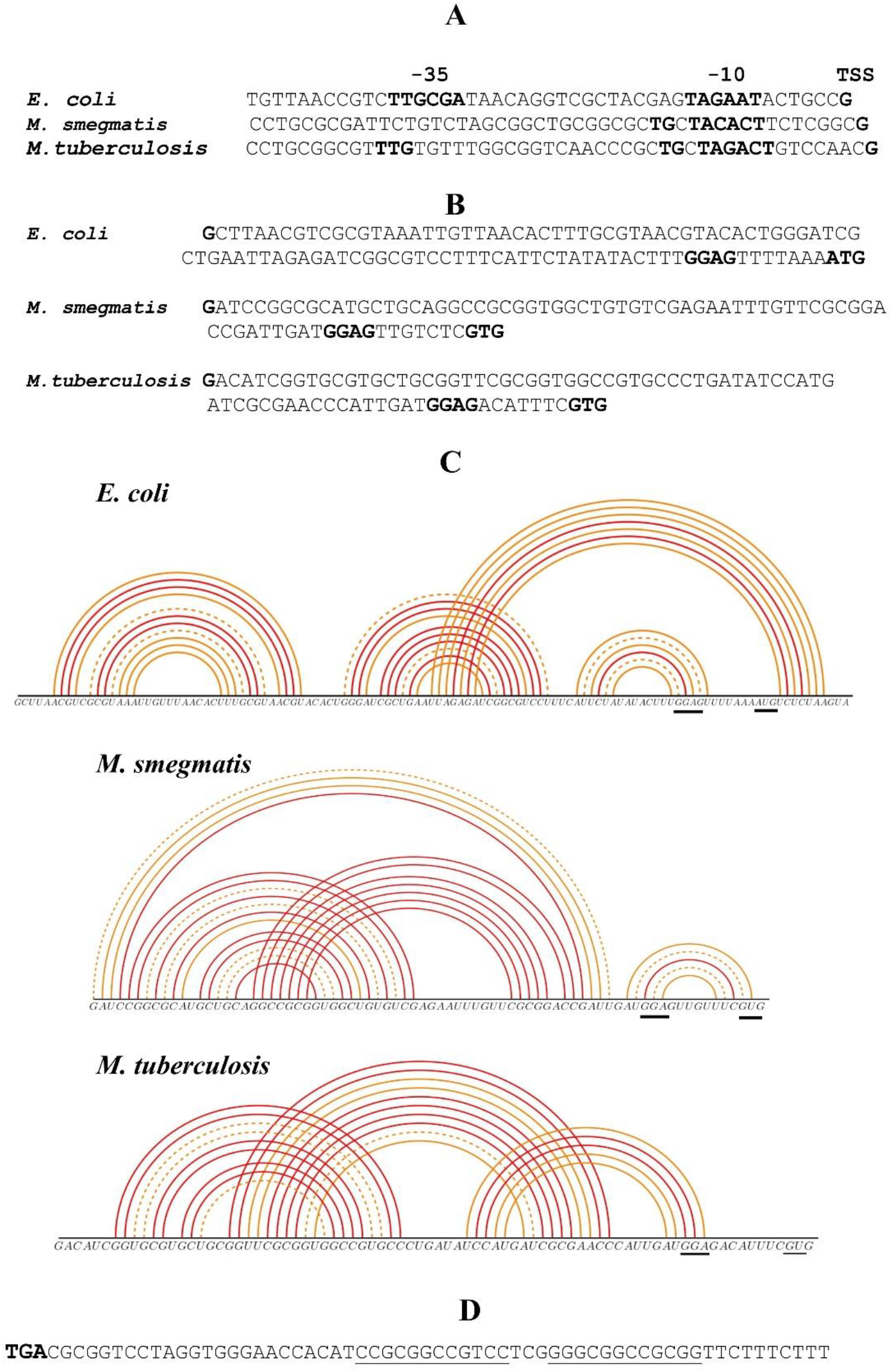
Regulatory elements in front of the *rpsO* coding region in *E. coli, M. smegmatis* and *M. tuberculosis*. A, B. Comparison of the core *rpsO* promoters (A) and 5’UTRs (B). TSS, −10, −35 promoter regions (A) as well as the initiator codon and the Shine-Dalgarno element (B) are in bold. (C) 5’UTRs of *E. coli, Msm* and *Mtb rpsO* mRNAs form pseudoknots (according to ref. 37). (D) Intrinsic transcription terminator of the *Msm rpsO* gene; complementary sequences forming a hairpin structure are underlined.

It should be underlined, the 5’ UTRs of *E. coli rpsO* as well as both mycobacterial *rpsO* 5’UTRs fold into pseudoknots (Fig. 1C). The main difference is that in *E. coli*, the pseudoknot includes the beginning of the coding sequence, while in *Msm* and *Mtb*, it is likely formed upstream of the start codon, according to the McGenus algorithm (37). In *E. coli*, the pseudoknot in the *rpsO* TIR plays a key role as an autogenous operator, providing the S15-mediated ribosome entrapment in a non-productive initiation complex (9, 13). It would be interesting to ascertain whether the mycobacterial *rpsO* pseudoknots could act in a similar way. Finally, the *Msm rpsO* operon bears an *E. coli*-like intrinsic transcriptional terminator represented by a strong hairpin-loop structure followed by a U-rich stretch (Fig. 1D), which is very convenient for constructing the plasmid for the ectopic expression of *Msm rpsO* in *E. coli*. Thus, at first sight, one can expect that the regulatory *rpsO* regions of *Msm* or *Mtb*, including the promoter and TIR, can be recognized by transcription/translation machineries of *E. coli*. However, further experiments showed that the actual situation appeared to be different.

### The *rpsO* promoter from *M. smegmatis* is inoperative in *E. coli*

Followed the strategy described above, we constructed the reporter *Msm rpsO’-’lacZ* under the control of the *Msm rpsO* promoter and TIR, first on the plasmid and then integrated it into the chromosome of ENS0, the *E. coli* Lac^-^ strain (Fig. 2A). Although we managed to obtain the Lac^+^ phenotype resulting from homologous recombination, the β-galactosidase assay showed a low expression output insufficient for statistically reliable measurements. In attempting to increase the expression level, we changed the *Msm rpsO* promoter for the promoter of *E. coli rpsO* while preserving the *Msm rpsO* 5’UTR intactness, which is indispensable for studying the *Msm rpsO* autogenous regulation. The resulting construct showed a ca 10-fold higher expression level (Fig. 2B), thus allowing us to evaluate an impact of *Msm* S15 *in trans* on the *Msm rpsO*-*lacZ* reporter.

**Figure 2.**
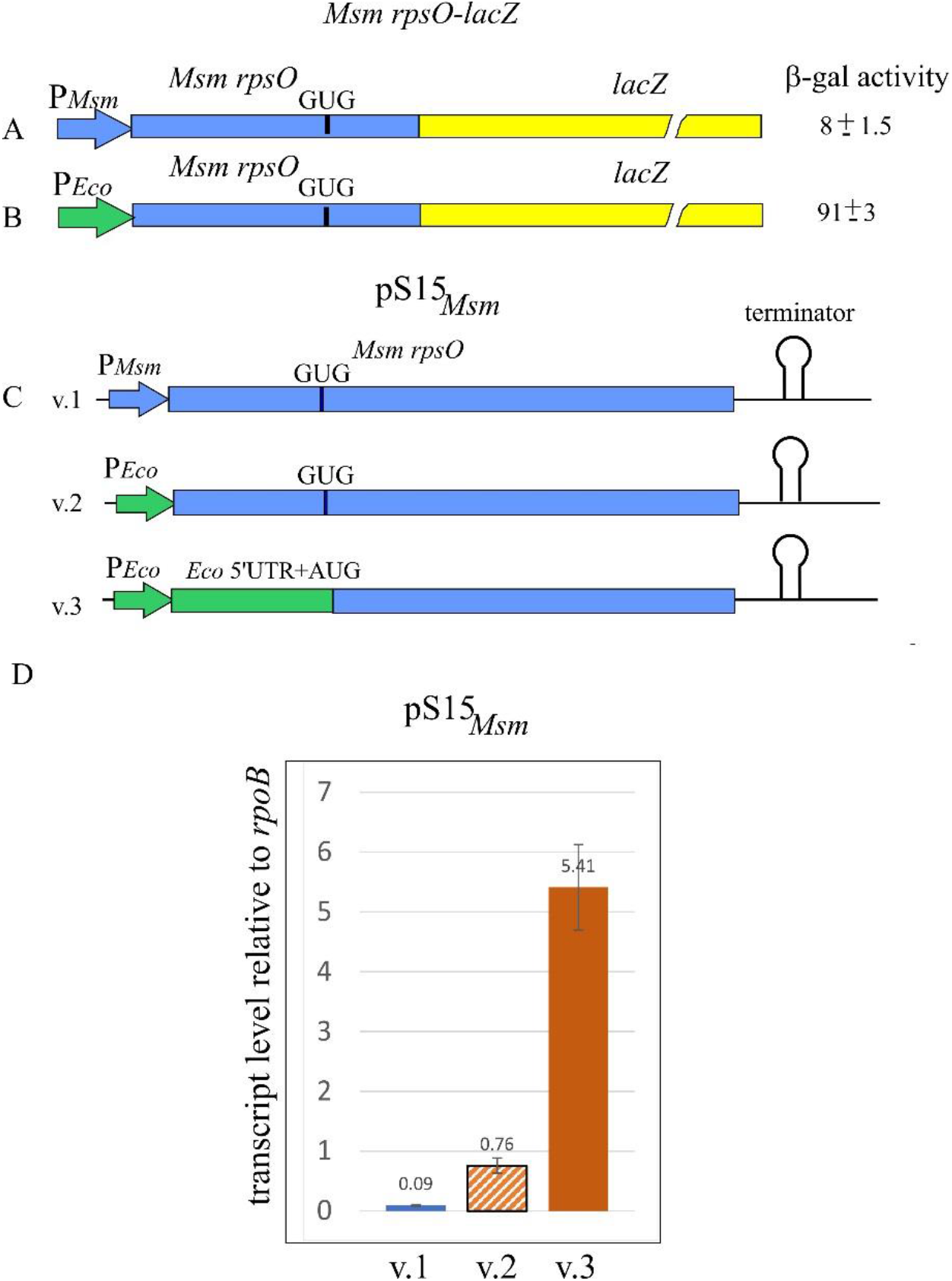
Constructs for studies of the *Msm rpsO* autoregulation by using *E. coli* as a surrogate host. A. *Msm rpsO’-’lacZ* fusion under the *Msm rpsO* promoter. B. *Msm rpsO’-’lacZ* fusion under the *E. coli rpsO* promoter. Corresponding β-galactosidase activities are indicated. C. Three versions of pS15_*Msm*_ plasmid (v.1, v.2, v.3) for expression of *Msm* S15 *in trans*. D. *Msm rpsO* transcript levels in *E. coli* cells bearing three versions of pS15_*Msm*_. The results of RT-qPCR analysis with the internal control *rpoB*. Transcript amounts relative to *rpoB* are indicated above the bars.

To provide the *Msm rpsO* ectopic expression, we constructed three versions of the plasmid pS15_*Msm*_ (Fig. 2C). First, using pACYC184, we cloned the whole gene *Msm rpsO* with its native flanks including the promoter, 5’UTR and the transcription terminator (version 1); in a version 2, the *Msm rpsO* promoter was changed for the *E. coli* counterpart, with the *Msm rpsO* 5’UTR remaining intact; finally, in a version 3, we replaced not only the *Msm rpsO* promoter, but also the 5’UTR and the initiator GUG with the *E. coli rpsO* promoter, 5’UTR and the initiator AUG (Fig. 2C).

Efficiency of the *Msm rpsO* gene expression from the constructed plasmids (pS15_*Msm*_ versions 1, 2 and 3) was evaluated by measuring the *Msm rpsO* transcript level in *E. coli* cells harboring the plasmids by using RT-qPCR, with the *rpoB* transcript serving as an internal standard (Fig. 2D). The highest level of the *Msm rpsO* transcript was found in cells bearing pS15_*Msm*_ v.3 where the synthesis of *Msm* S15 was driven by the regulatory regions of *E. coli rpsO*. The plasmid pS15_*Msm*_ v.1 (*Msm rpsO* promoter, *Msm rpsO* 5’UTR) showed the lowest transcript yield (Fig, 2D), in line with the low expression output of the *Msm rpsO’-’lacZ* reporter. Importantly, the change of only the *Msm rpsO* promoter for the *E. coli* counterpart (pS15_*Msm*_ v.2) significantly increased the transcript level, suggesting that the mycobacterial *rpsO* promoter is inoperative in *E. coli*.

It is of interest, not only the promoter region but also the 5’UTR structure had a great impact on the transcription output. A comparison of the *Msm rpsO* transcript levels for cells bearing pS15_*Msm*_v.2 and pS15_*Msm*_v.3 revealed a 7-fold increase, indicating that transcription and hence overall expression of the GC-rich *Msm rpsO* coding sequence becomes more efficient with the *E. coli* 5’UTR, despite the presence of the same *E. coli rpsO* promoter (Fig. 2D). We suppose that the cognate *E. coli rpsO* TIR provides much more effective ribosome loading during translation initiation, thereby ensuring efficient transcription-translation coupling necessary for the synthesis of a stable transcript (38, 39 and references therein). Furthermore, it has been shown that the r-protein S1 plays a key role in recognition and binding of mRNA 5’-UTRs by the *E. coli* 30S ribosomal subunit during initiation complex formation (40, 41), including the structured *rpsO* mRNA 5’UTR able to form a pseudoknot (42). It should be noted that an ability of S1 to unfold RNAs forming pseudoknots inversely correlates with their structural stability (43). This may suggest that the *Msm rpsO* 5’UTR able to form a stable pseudoknot (Fig. 1C) represents an arduous target for *E. coli* S1. In addition, it has been demonstrated that the *E. coli* S1 capacity to recognize 5’UTRs of high GC-mRNAs is limited, so that the high GC content of heterologous mRNAs presents a significant challenge to *E. coli* ribosomes when initiating translation (44). This was supported by the directed evolution of S1, resulting in selection of S1 mutants capable of enhancing translation of GC-rich mRNAs by *E. coli* ribosomes (44). Thus, the limited ability of *E. coli* S1 to recognize and bind GC-rich sequences within 5’UTRs is, most likely, the main reason behind the low expression level of the *Msm rpsO* mRNA in *E. coli*.

### The *Msm rpsO* gene is feedback regulated at the translation level

Based on the above observations (Fig. 2D), the plasmid pS15_*Msm*_v.3 was chosen for subsequent studies of the *Msm rpsO* autogenous control. The *E. coli* cells bearing the *Msm rpsO’-‘lacZ* reporter under the control of the *E. coli rpsO* promoter (Fig. 2B) were transformed with pS15_*Msm*_v.3 or with an empty vector. The resulting transformants were grown to the exponential phase (OD_600_∼0.4-0.5), and the corresponding β-galactosidase levels were measured. Although the expression of the reporter was not high, the use of 5 or more biological replicates allowed us to obtain statistically reliable results which revealed ca 6-fold repression in the presence of pS15_*Msm*_v.3, thus clearly indicating the feedback regulation of the *Msm rpsO* mRNA (Fig. 3A). The repression level was about the same as for *E. coli rpsO* in the presence of pS15_*Eco*_, even though the expression of the *Eco rpsO-lacZ* reporter was incomparably higher (Fig. 3B). Intriguingly, pS15_*Eco*_ was also able to inhibit the expression of the *Msm rpsO’-’lacZ* reporter, with the repression level being a bit lower (Fig. 3A). At the same time, pS15_*Msm*_v.3 had only marginal impact on the expression of *Eco rpsO’-’lacZ* (Fig. 3B), indicating that despite the high homology level of the S15 proteins (Fig. 3C), S15_*Msm*_ is not capable of recognizing the *E. coli rpsO* operator, whereas S15_*Eco*_ has an ability to bind the heterological *rpsO* 5’UTR and to inhibit translation, presumably due to its higher RNA-binding plasticity. The results obtained indicate that the *Msm rpsO* mRNA is feedback regulated at the translation initiation level, just as the *E. coli rpsO*. Most likely, *Msm* S15 recognizes some structural features of the *Msm* 5’UTR structure, thereby stabilizing the pseudoknot and preventing ribosome loading. The mechanistic details of the S15-mediated autogenous control in mycobacteria represent an interesting issue to be addressed in future.

**Figure 3.**
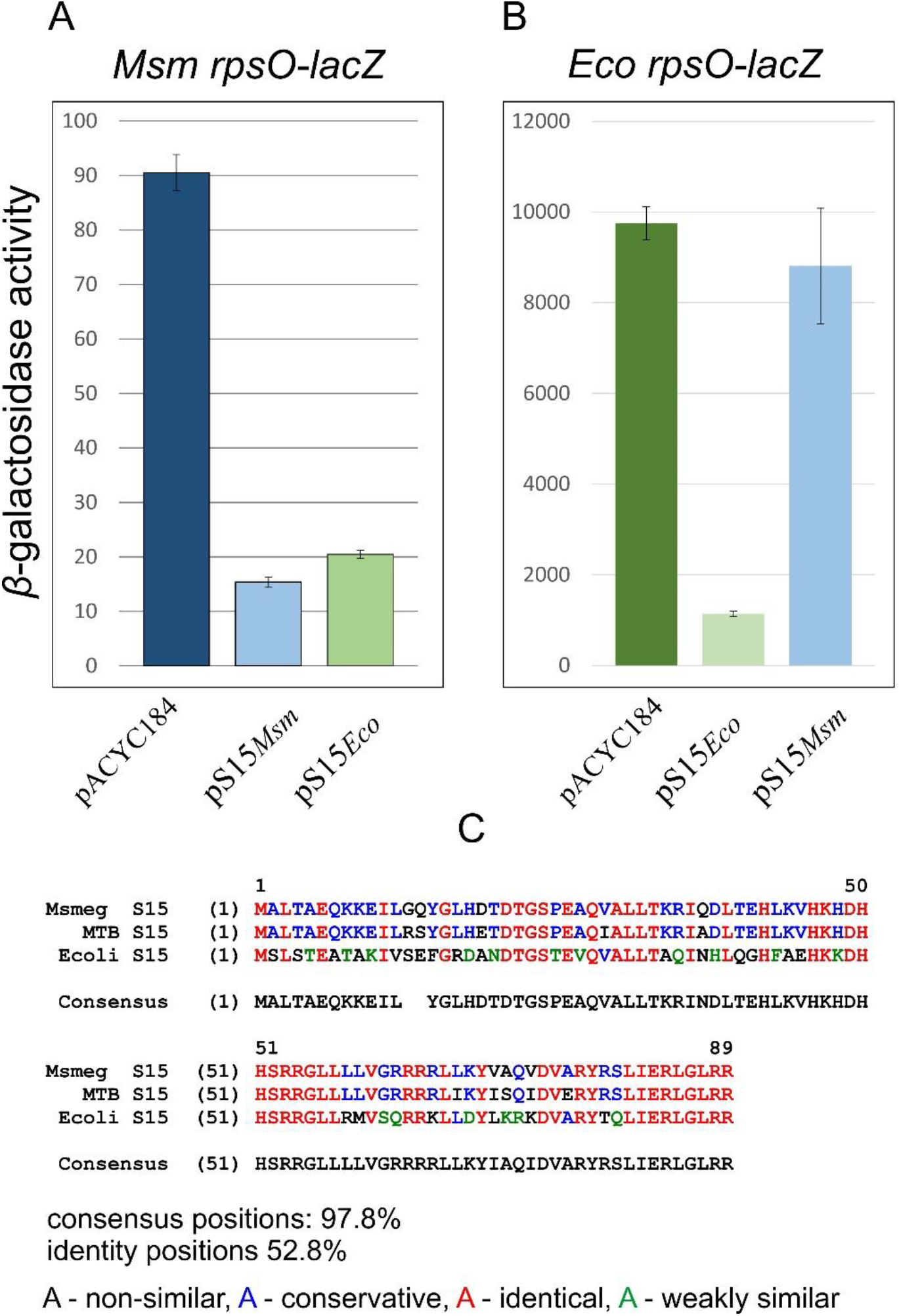
The *Msm rpsO* gene is feedback regulated at the translation level, like *E. coli rpsO*. A. Inhibition of the *Msm rpsO-lacZ* reporter expression in the presence of S15_*Msm*_ and S15_*Eco*_ *in trans*. B. Autogenous regulation of the *E. coli rpsO-lacZ* reporter: S15_*Eco*_ in trans inhibits expression, while S15_*Msm*_ has only marginal effect. C. Homology of S15 from *E. coli, Msm* and *Mtb*.

### Creation of the cognate system for studying the autogenous control of r-protein synthesis in mycobacteria

The low expression output of the *Msm rpsO’-’lacZ* reporter revealed obvious limitations of using *E. coli* as a surrogate host to study the autogenous control of mycobacterial r-proteins. Indeed, to make constructs able to provide a measurable efficiency of the reporter integrated in the chromosome, we had to change the mycobacterial *rpsO* promoter for the *E. coli* counterpart. Further, to provide efficient expression of *Msm* S15 in trans, we had to substitute the regulatory region including the promoter, 5’UTR and the start codon for *E. coli* determinants. In addition, S15 is a small protein (89 amino acid residues), so that expression of the *Msm rpsO* short coding region in *E. coli* appeared to be not a very difficult task for transcription/translation machineries of *E. coli* even though they had been adapted to a lower GC content. It is reasonable to suspect that the mycobacterial mRNAs encoding longer r-proteins (like RpsA, RpsB) will bring much more problems, making the use of *E. coli* as a surrogate host unpromising for future studies. Thus, it is vital to develop the authentic system for studies the r-protein-mediated control in mycobacteria, and *M. smegmatis* represents the best proxy for such experiments (22).

Both integrative (to be inserted into mycobacterial chromosome) and replicative (for ectopic expression of genes under study) plasmids for creating the *Msm*-based reporter system were reported (23) and widely used. The integrated plasmid pMV306hsp was initially derived from a replicative vector pMV261 by replacing the mycobacterial replication origin (*oriM*) with a DNA fragment comprising the attachment site *attP* and the integrase gene *int* from the mycobacteriophage L5, which provided a site-specific integration into the chromosomal *attB* site (23). In addition, this integrative vector carries the *hsp*60 promoter and the *rrnB* terminator to provide cloning and expression of different genes as a single copy integrated into the chromosome. We modified pMV306hsp by replacing the region comprising the *hsp* promoter with the *Msm rpsO-egfp* reporter bearing the *Msm rpsO* core promoter and 5’UTR in front of the eGFP coding sequence, so that transcription of the reporter gene would be governed by the *rpsO* core promoter and terminate at the *rrnB* terminator (Fig. 4A). The newly created integrative plasmid pMV*rpsO-egfp* (version1) was checked by sequencing and used to transform *M. smegmatis* cells by electroporation resulting in kanamycin-resistant transformants.

**Figure 4.**
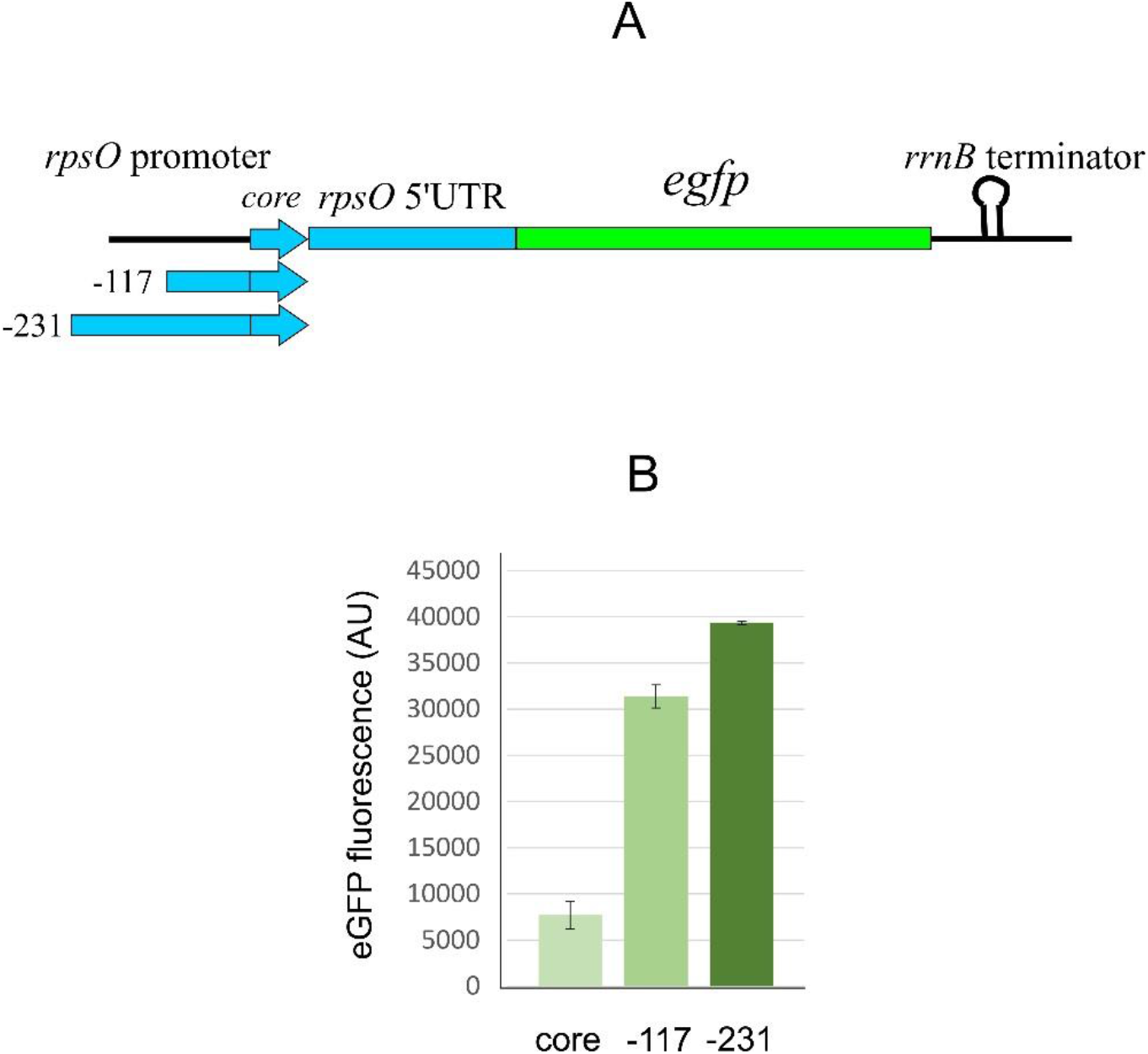
Expression level of the chromosomally integrated *Msm rpsO-egfp* reporter depends on the 5’-extension of the *Msm rpsO* promoter. A. *Msm rpsO-egfp* reporters within the *Msm* chromosome. B. Results of fluorescent measurements of protein lysates obtained from exponential *Msm* cells bearing *rpsO-egfp* reporters governed by the *rpsO* promoters with different 5’ extensions; positions relative TSS are indicated, with “core” corresponding to the 5’ edge position −47.

### The *Msm rpsO* core promoter requires an upstream region to enhance the transcription yield

To evaluate the efficiency of the chromosomally integrated fluorescent reporter, the *Msm* cells (kan^r^) were harvested at the exponential phase (OD_600_∼0.7-0.8), and then disintegrated for preparing protein lysates to use for measuring the fluorescence of the reporter. The fluorescence appeared unexpectedly low despite that, as a rule, ribosomal core promoters, at least in *E. coli*, are effective (e.g., the *rpsO* promoter in a fusion *Eco*_*rpsO-lacZ*, see Fig. 3B). Analysis of the published data revealed that, in contrast to *E. coli*, mycobacterial core promoters (including only −10 and −35 promoter regions) may be inefficient requiring 5’extensons to augment their strength (28, 45, 46). In particular, the core *Msm rrnB* promoter (which a priori should be one of the strongest in bacterial cells) remained relatively weak unless the upstream region was significantly extended (28). To test whether it is also the case of the *rpsO* gene, we extended the *rpsO* promoter sequence (the initial 5’ edge was at the position −47 from TSS) to obtain the 5’ extended variants: version 2 (−117) and version 3 (−231), and then created the *Msm* cells bearing the corresponding *rpsO-egfp* reporters in the chromosome (Fig. 4 A). The fluorescence measurements revealed that the extended promoter variants gave the increased yield of eGFP (Fig. 4B). The same was previously shown for the *Msm rrnB* promoter (28); in addition, the promoters of the sRNAs Ms1 (45) and MrsI (46) were also reported to be active in 5’ extended versions. The exact mechanism for the enhancement of transcription efficiency upon the promoter extending has not yet been clarified. One of the reasonable explanations is the existence of the upstream binding sites for as-yet unknown transcription factors acting as activators (28, 45). Based on the experimental observations, we used the *Msm rpsO*-*egfp* fusion bearing the −231 extension for further experiments.

### Generation of the novel replicative shuttle vector, pAMYC

To provide the ectopic expression of the *Msm rpsO* gene necessary to study the S15 −mediated effect on the efficiency of the *rpsO-egfp* reporter, we created a novel shuttle replicative vector, pAMYC, by transferring the region comprising the mycobacterial replication origin from pMV261 to pACYC184 (Fig. 5). The vector pMV261 itself is not applicable in our case because it bears the same kanamycin-resistance marker as an integrative pMV306 used for incorporation of the *rpsO-egfp* fusion into the *Msm* chromosome. Electroporation of a novel shuttle plasmid pAMYC into *Msm* cells yielded the chloramphenicol resistant transformants, indicating its suitability for ectopic expression of different mycobacterial genes (Fig. 5).

**Figure 5.**
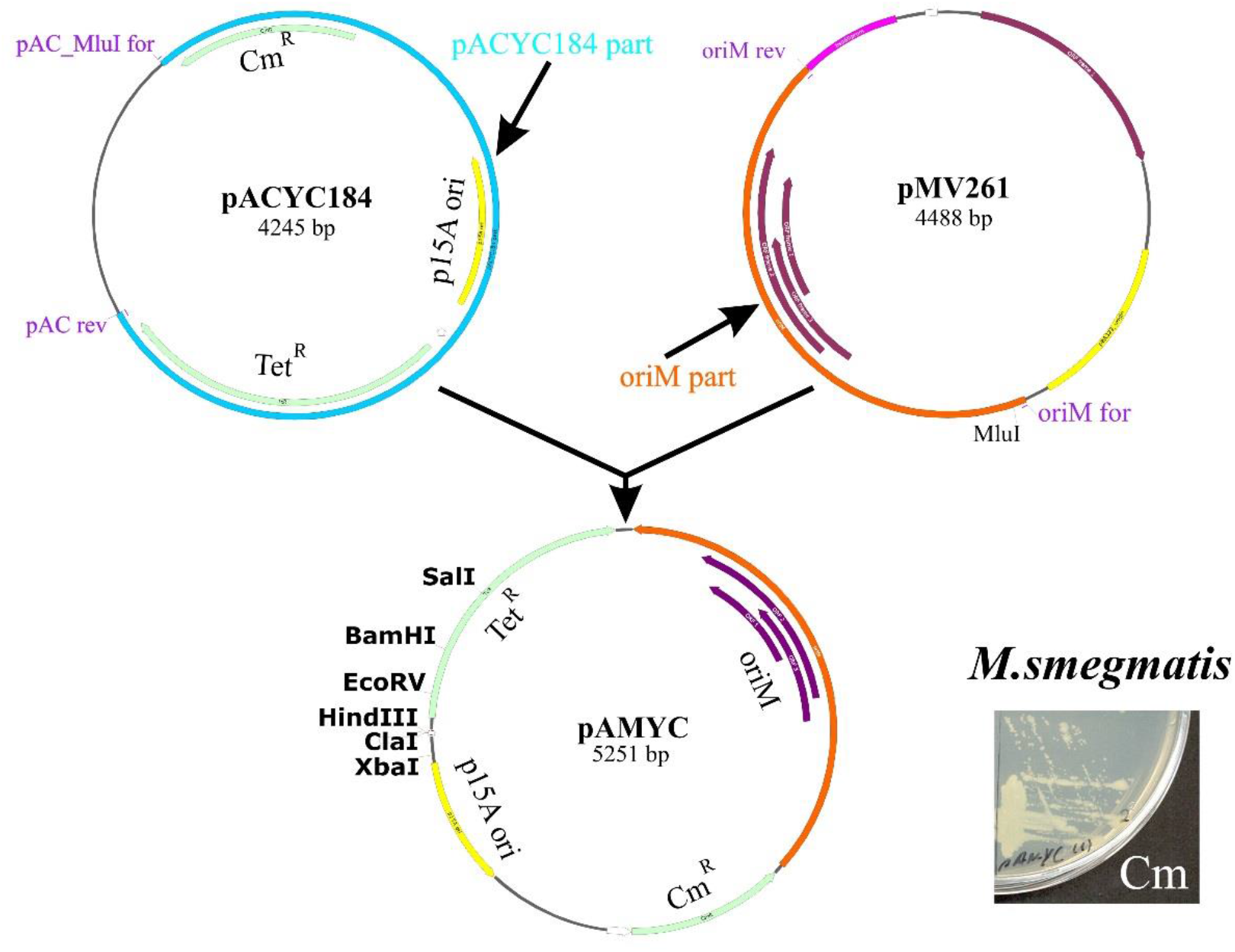
The scheme for creating a novel replicative shuttle vector pAMYC on the basis of pACYC184. Transformation of *M. smegmatis* by pAMYC results in appearance of chloramphenicol resistant cells on LB-Cm plates.

### Mycobacterial *rpsO* expression is feedback regulated at the translation level

To obtain the pAMYC derivative providing the synthesis of *Msm* S15 in trans, we cloned the *Msm rpsO* gene bearing the 5’-extended promoter variant (−231) and its own intrinsic terminator. The resulting plasmid pAMS15_*Msm*_ was sequenced and used to transform *Msm* cells bearing the chromosomal *rpsO-egfp* fusion under the same 5’-extended (−231) *rpsO* promoter. An empty pAMYC served as a control. Exponential *Msm* cells were harvested and disintegrated to prepare protein lysates where the eGFP fluorescence was measured. The results clearly showed the reduced fluorescence in cells bearing pAMS15_*Msm*_ when compared to the control cells bearing an empty pAMYC (Fig. 6A), thus indicating that S15 *in trans* down-regulates the reporter expression. This supports the results obtained with *E. coli* as a surrogate host.

**Figure 6.**
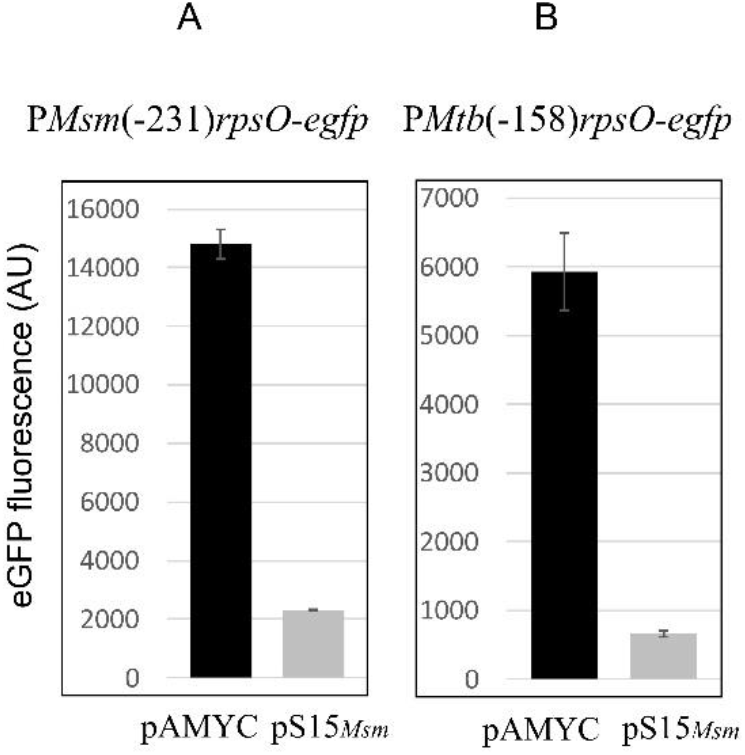
The *rpsO* genes of *M. smegmatis* and *M. tuberculosis* are feedback regulated in vivo at the translation level. Repression of the *Msm rpsO-egfp* (A) and *Mtb rpsO-egfp* (B) expression in the presence of the pAMYC derivative expressing *Msm* S15. Results of fluorescent measurements.

Further, we created analogous chromosomally integrated reporter where the eGFP expression was under the control of the extended (−158) promoter and 5’UTR of the *Mtb rpsO* gene. The fluorescence was measured for the *Msm* cells bearing the pAMYC derivative expressing *Msm* S15 (which has 88.8% of identity with the *Mtb* S15) and an empty vector as a control. The results clearly showed the repression of the *Mtb rpsO-egfp* expression by *Msm* S15 in trans (Fig. 6B). Taken together, the results allow us to conclude that mycobacterial *rpsO* expression is feedback regulated at the translation level, and that the mechanism likely resembles that of *E. coli*. Most probably, acting as a repressor, S15 binds the 5’UTR folded in a pseudoknot, thereby stabilizing its structure that impedes ribosome loading to initiate translation. In this regard, the mechanism of the *Msm rpsO* repression seems to differ from the “entrapment” of the 30S subunit in a nonproductive complex observed in *E. coli*. Future studies may shed light on the exact mechanism of the autogenous control of the mycobacterial *rpsO* gene.

### Concluding remarks

In this work, we provide evidence that the autogenous control of r-protein synthesis at the translation level is functioning in mycobacteria. We developed a reporter system based on *M. smegmatis* which is suitable to study the regulation of mycobacterial genes encoding r-proteins. The use of *E. coli* as a surrogate host for this purpose was found unpromising because of a low efficiency of *E. coli* transcription/translation machinery in expression of mycobacterial genes. To study the regulation of the *rpsO* gene encoding r-protein S15, we obtained the *Msm* cells (kan^r^) bearing reporters *Msm_rpsO-egfp* and *Mtb_rpsO-egfp* in the chromosome and measured their activity in the presence of the *Msm* S15 expressed from a new replicative shuttle plasmid pAMYC (Cm^r^) vs an empty vector. The inhibition of the reporter expression by *Msm* S15 in trans clearly indicated the autogenous control of *rpsO* expression in mycobacteria.

Like the *rrnB* (28) or Ms1 (45) core promoters, the *rpsO* core promoter of *M. smegmatis* appeared rather weak requiring the upstream sequences for stimulating its activity. As the mycobacterial *rpsO* genes are transcribed from a single promoter, according to the RNAseq data (47, 48), the stimulation by the upstream sequences cannot be due to the activity of additional upstream promoters but rather may be explained by the action of as-yet-unknown transcription factors the binding sites of which are situated within the upstream sequence (upstream activating elements).

We believe that the proposed reporter system is applicable for studies of different regulatory circuits involving proteins or sRNAs as mRNA targeting *trans* factors in mycobacteria and, most likely, in various other species of Actinobacteria.

## MATERIALS and METHODS

### Strains and plasmids

Strains and plasmids used in this study are listed in Table 1. *Mycobacterium smegmatis* mc^2^155 was provided by Prof. A. S. Kaprelyants (Bach Institute of Biochemistry, Moscow). Isolation of *M. smegmatis* genomic DNAs was performed according to Belisle et al. (49). *M. tuberculosis* genomic DNA was a kind gift of Dr. E. Salina (Bach Institute of Biochemistry, Moscow). For experiments with *E. coli* as a surrogate host, plasmids pS15_*Msm*_ (versions 1, 2, 3), derivatives of the pACYC184 cloning vector, were constructed to express *in trans* the *rpsO* gene from *M. smegmatis* (*Msm*). The plasmid pEMsm_rpsO-lacZ, a derivative of pEMBLΔ46 (24) bearing the *Msm rpsO’-’lacZ* fusion, was used for transferring this reporter onto the chromosome of ENS0 by homologous recombination. For the *M. smegmatis* expression system, the derivatives of the pMV306 integrative plasmid bearing the kanamycin-resistance marker (23) and a novel replicative shuttle-vector pAMYC (providing chloramphenicol resistance) were created (see below).

**Table 1.**
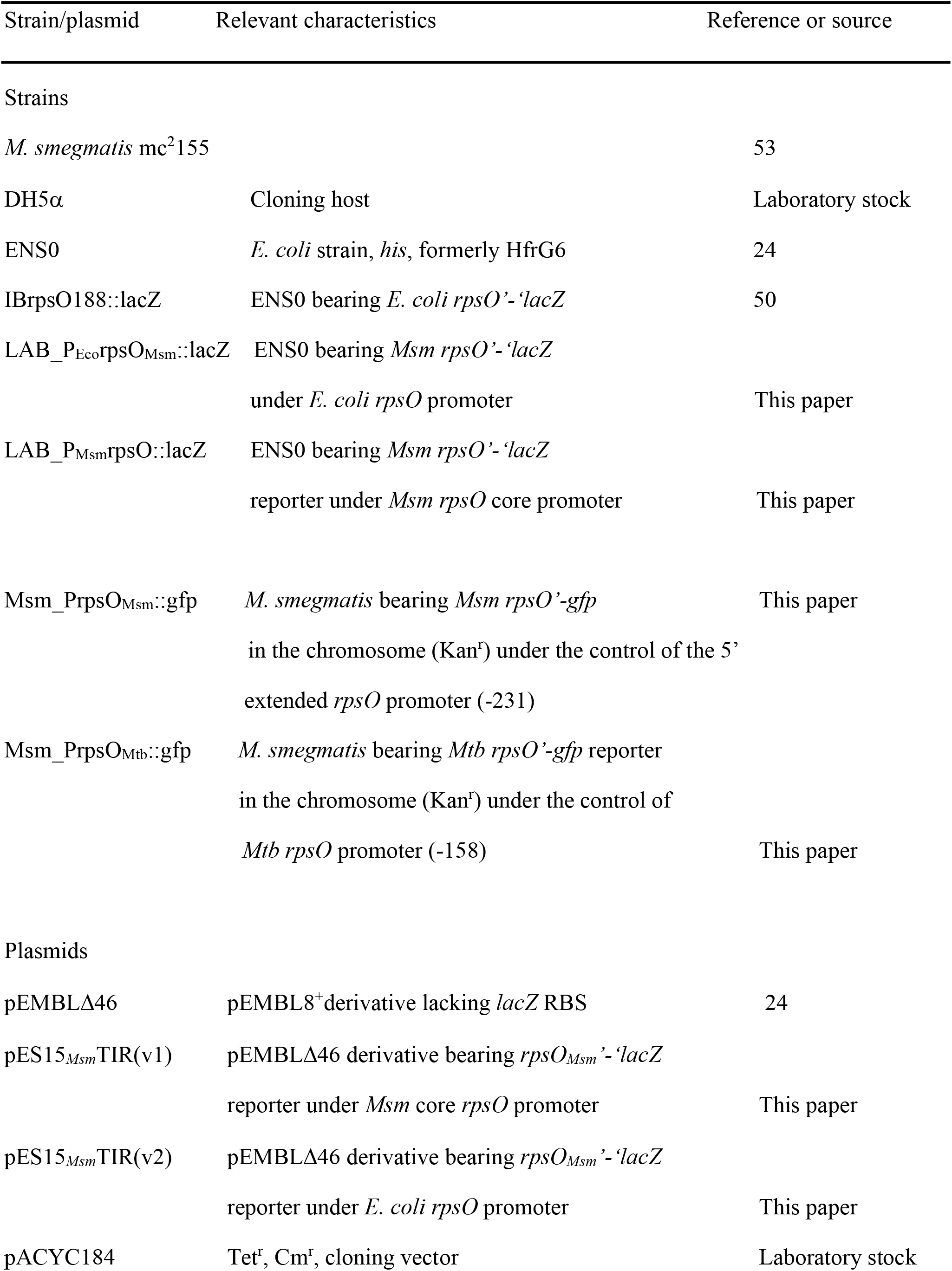
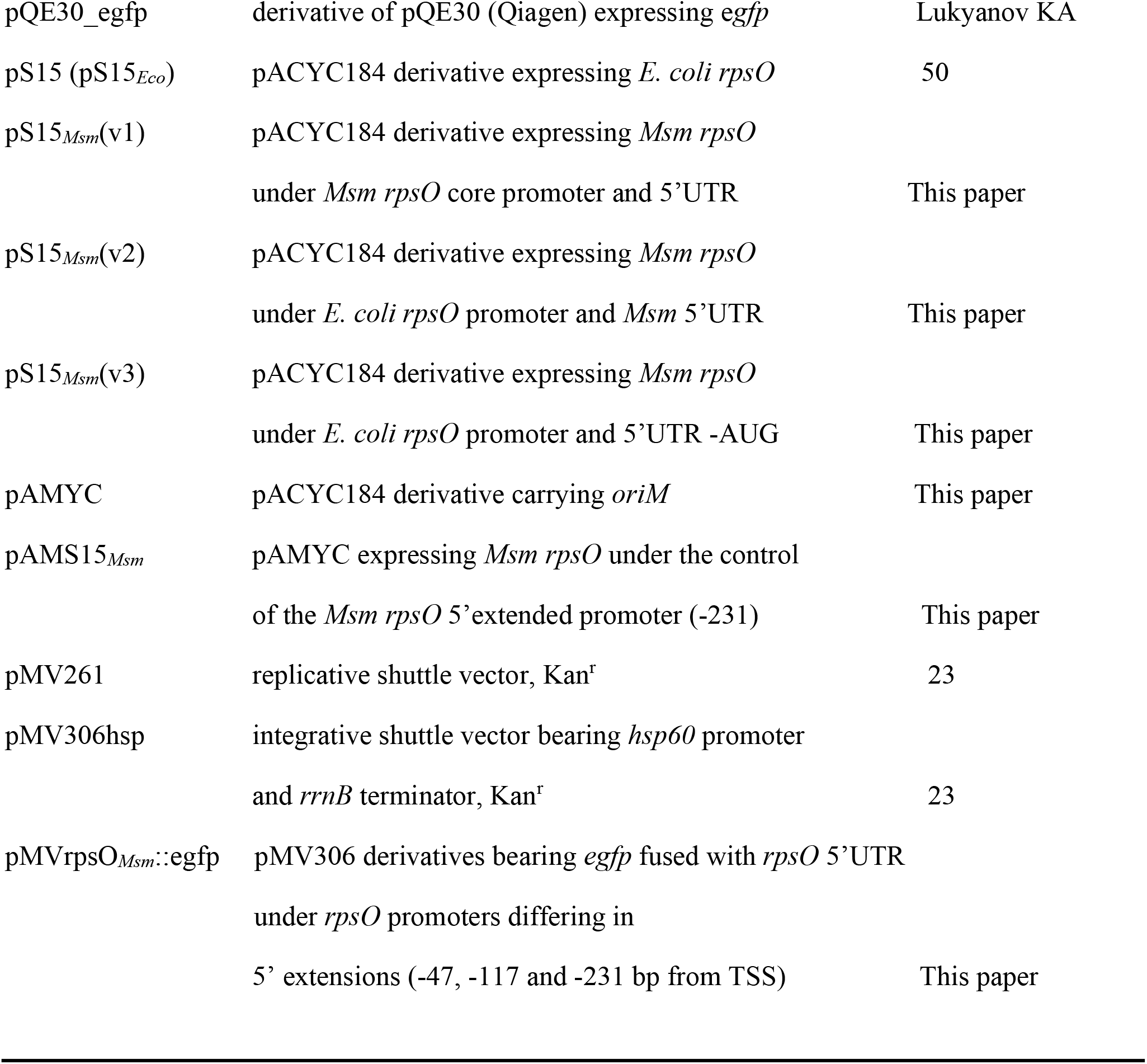
Strains and plasmids used in this study.

### Construction of expression plasmids for using in *E. coli* as a surrogate host

To generate pS15_*Msm*_ (v.1), the *rpsO* gene flanked with its own promoter and terminator sequences was amplified by PCR on *Msm* genomic DNA with primers Msm-rpsO-for 5’-ATC***GGATCC***GCACGATCCTGC and Msm-rpsO-rev 5’-ACT***AAGCTT***GCATGTCCGCAGAC. Forward and reverse primers comprised BamHI (for) and HindIII (rev) sites (bold italicized) for subsequent cloning in pACYC184. To create pS15_*Msm*_ (v.2), we replaced the *Msm rpsO* promoter with the *E. coli rpsO* promoter by using a two-step PCR technique. First, we obtained two PCR fragments; one of them was amplified on pS15_*Eco*_ (a pACYC184 derivative bearing the *E. coli rpsO* gene flanked with its native promoter and terminator, see 50), using the forward primer corresponding to the pACYC184 sequence including BamHI site (pACYC184-for 5’-CGATGCGTCCGGCGTAGA***GGATCC***) and the reverse primer (PrpsOmix-rev) comprising a sequence complementary both to the *E. coli rpsO* promoter/discriminator region and the beginning of the *Msm rpsO* transcript (5’-<u>CATGCGCCGGATC</u>GGCAGTATTCTACTC, with the *Msm* sequence underlined). Another PCR fragment was amplified on pS15_*Msm*_ (v.1) with primers PrpsOmix-for, a complement of the PrpsOmix-rev (5’-GAG**TAGAAT**ACTGCC<u>GATCCGGCGCATG</u> where the *Msm* sequence underlined, *Eco_rpsO* −10 promoter in bold), and Msm-rpsO-rev described above. At the second step, the two PCR fragments were mixed and amplified with external primers pACYC184-for and Msm-rpsO-rev; the resulting product was treated with BamHI and HindIII and cloned in pACYC184/BamHI, HindIII.

Lastly, to create pS15_*Msm*_ (v.3), we substituted not only the *Msm rpsO* promoter but also 5’-UTR and the start codon GUG for corresponding *E. coli* sequences. At the first step, two PCR fragments were obtained: one amplified on pS15_*Eco*_ with primers pACYC184-for (see above) and Eco-rpsOTIR-rev (5’-<u>CGGCGGTAAGCGC</u>CATTTTAAAACTCCAAAG, where the *Msm* sequence is underlined), and another one amplified on pS15_*Msm*_ (v.1) with primers Eco-rpsOTIR-for (5’-CTTT**GGAG**TTTTAAA**ATG**<u>GCGCTTACCGCCG</u>, where *E. coli* SD-sequence and AUG start codon in bold and the *Msm rpsO* sequence underlined) and Msm-rpsO-rev. At the second step, the two PCR fragments were mixed and amplified in the presence of pACYC184-for and Msm-rpsO-rev, the resulting product was cloned in pACYC184 as described above. All three versions of pS15_*Msm*_ were sequenced and used for further experiments.

### Quantification of the in vivo transcripts by RT-qPCR with an internal standard

The efficiency of the *Msm rpsO* gene expression in *E. coli* was evaluated by measuring the *Msm rpsO* transcript levels in cells bearing plasmids pS15_*Msm*_ (versions 1, 2, 3) or the parental empty vector pACYC184 as a control. Total RNA was isolated by using the RNeasy mini kit (Qiagen) according to recommendations of the manufacturer. Strains (including the control bearing an empty vector) were grown in LB medium at 37°C with vigorous shaking. At an optical density at 600 nm (OD_600_) of ∼0.4, 2-ml aliquots of cell cultures were withdrawn and mixed with 4 ml RNAprotect bacterial reagent. Further, total RNA was extracted; during extraction RNase-free DNase was added to the columns for 15 min to eliminate DNA contaminations in RNA samples; after elution RNA concentrations were estimated by measuring the OD_260_. Reverse transcription (RT) was performed with AMV reverse transcriptase (Promega) in final volume 20 μl for 1h at 42°C on 1 μg of total RNA in the presence of two reverse primers (1 μl of 5 μM solution each) corresponding to the coding part of *Msm rpsO* (Msm-rpsOcod-rev 5’-CGAGTGGTGATCGTGCTTGTGC) and to the reference gene *rpoB* (rpoB-rev: 5’-CGGATTTGACATTCCTGGACGTC). Real-time PCR (qPCR) was run with the use of LightCycler 96 (Roche); each 25 μl reaction contained 2 μl RT mix, 5μl 5x qPCRmix HS SYBR (Evrogen), forward primers corresponding to the beginning of transcripts (Msm-rpsOtr-for 5’-GTGGCTGTGTCGAGAATTTGTTCG for pS15_*Msm*_ v.1 and 2, Eco-rpsOtr-for 5’-GTAACGTACACTGGGATCGCTG for pS15_*Msm*_ v.3, rpoB-for 5’-ACGTCCACAAGTTCTGGATGTACC) and reverse primers used for RT (1 μl of 5μM solution each). Two independently isolated preparations of total RNA for each of 4 strains were used for RT, and three technical replicates for each qPCR reaction were run simultaneously. Control qPCR reactions without RT were done to exclude DNA contaminations in RNA preparations. LinRegPCR software was used to quantify transcript amounts relatively the reference transcript *rpoB*.

### Construction of the *Msm*_*rpsO’-’lacZ* fusions integrated into *E. coli* chromosome

The *Msm_rpsO’-’lacZ* chromosomal fusions were generated as previously described for different fusions related to the r-protein operons from γ-proteobacteria (see e.g. 25, 26). For the fusion under the control of the *Msm rpsO*-promoter, a DNA fragment was amplified on pS15_Msm_v1 with primers Msm-rpsO-for (see above) and Msm_TIR_rpsOrev 5’-GG*AAGCTT*TGGCCCAGGATCTC. The forward primer comprised BamHI site, the reverse primers -HindIII (italicized in the sequence). The resulting fragment was cloned in pEMBLΔ46/BamHI, HindIII in frame with the *lacZ* open reading frame, then the correct construct was transferred onto the chromosome of ENS0 (24) by homologous recombination followed by selecting the recombinant Lac^+^ strains on McConkey agar (24). To substitute the *Msm rpsO* promoter with the *E. coli* counterpart, the corresponding DNA fragment was amplified on pS15_Msm_v2 with primers pACYC184-for and Msm_TIR_rpsOrev (see above), cloned in pEMBLΔ46/BamHI, HindIII and then transferred onto the chromosome of ENS0 as described above.

### Cell growth and β-galactosidase assay

*E. coli* cells bearing the *Msm rpsO’-’lacZ* reporters and the plasmid expressing *Msm* S15 or the empty vector were grown at 37^°^C in Luria-Bertani (LB) medium supplemented with chloramphenicol (34 μg/ml), harvested in exponential phase at OD_600_ ∼ 0.4-0.5 and used for preparing clarified cell lysates essentially as described (25). Protein concentration in each fraction of soluble proteins was determined by Bradford assay (Bio-Rad). Specific ß-galactosidase activities in the same fractions were measured according to Miller (51) and expressed in nmol ONPG (*O*-nitrophenyl-β-D-galactopyranoside) hydrolyzed per minute per milligram of total soluble cell proteins.

### Creation of a novel *Escherichia coli*-mycobacteria shuttle vector pAMYC, a derivative of pACYC184

A 3328 bp-fragment of pACYC184 comprising genes for chloramphenicol (Cm) and tetracycline (Tet) resistance as well as a replication origin p15A (*oriE*) was PCR amplified by using Q5 High-Fidelity DNA Polymerase (New England Biolabs) with primers pACshtl-for (5’-TTC*ACGCGT*AGCACCAGGCG, MluI restriction site italicized) and pACshtl-rev (5’-CTCCGCAAGAATTGATTGGCTCC). Mycobacterial origin of replication (*oriM*) was amplified from the plasmid pMV261 (23) by using Q5 DNA Polymerase and primers oriM-for (5’-GCCTTTGAGTGAGCTGATACCG) and oriM-rev (5’-GATTTAAAGATCTGGTACCGCGGC), resulting in a 1976-bp PCR fragment. The PCR fragments (3328 and 1976 bp in length) were gel-purified, treated with MluI (MluI site in the *oriM*-fragment is located near the annealing site for oriM-for), phosphorylated at blunt ends by treating with T4-PNK (Fermentas), and then ligated by T4-DNA ligase (Fermentas) at room temperature. Ligation mix was used to transform DH-5α cells; plasmids were isolated from Cm^r^-transformants and used for electroporation of *M. smegmatis* cells (52). Cm-resistant colonies appeared on LB-Cm agar plates after 3 days of incubation at 37°C, indicating that a newly created plasmid (named as pAMYC) indeed works as a mycobacteria-*E. coli* shuttle vector and thus may be used for cloning and expression *in trans* of mycobacterial proteins (or sRNAs, depending on the task) to study their effect on expression of mycobacterial mRNA targets.

### Modification of the integrative plasmid pMV306hsp to provide insertion of the *rpsO-egfp* reporter construct into the chromosome of *M. smegmatis*

The integrative shuttle vector pMV306hsp (23) carries the *hsp*60 promoter and the *rrnB* terminator to provide cloning and expression of different genes as a single copy integrated into the chromosome. We modified this plasmid by deleting the region comprising the *hsp60* promoter and instead inserting the *rpsO*_*Msm*_-e*gfp* reporter in front of the *rrnB* terminator. To this end, we first treated pMV306hsp with endonucleases MluI (upstream of the *hsp* promoter) and HindIII (in front of the *rrnB* terminator), and then dephosphorylated the treated vector with TSAP (Thermosensitive Alkaline Phosphatase, Promega).

To generate inserts comprising the *egfp* reporter under the control of the *Msm rpsO* regulatory regions (including the promoter and 5’-UTR), we made fusions *rpsO-egfp* by the two-step PCR technique with overlapping primers. For the first fusion, we used the *Msm rpsO* core promoter 5’-end of which corresponded to the −47 position from the transcription start site (TSS). The *rpsO* part was amplified from the *M. smegmatis* genomic DNA by using Tersus Plus PCR kit (Evrogen) with primers (−47) rpsO-for (5’-CTA*ACGCGT*TCCTGCGCGATTCTG, MluI site italicized) and rpsO-egfp--rev (5’-CGCCCTTGCTCACCACGAAACAACTCCA). The *egfp* part was amplified from the plasmid pQE30-egfp (Table 1) with primers rpsO-egfp-for (complementary to the rpsO-egfp-rev) and pQEegfp-rev (5’-GGAGTCCAAGCTCAGCTAATTAAGC, located downstream from HindIII site of pQE30-egfp). At the second step, the two PCR products were mixed and amplified with the external primers (−47) rpsO-for and pQE30egfp-rev, the resulting product was cleaned from 2% agarose gel by Cleanup Standard Kit (Evrogen), digested by MluI and HindIII and ligated into pMV306/MluI, HindIII. The reporter constructs bearing 5’-extended *rpsO* promoters were created in a similar way with the primers (−117) rpsO-for (5’-TCT*ACGCGT*AGGAGAAGTTCGATTC) and (−231) rpsO-for (5’-TGA*ACGCGT*AATCCGACGTTCTC), other primers were the same as described above.

The *rpsO-egfp* reporter construct bearing the 5’UTR and the *rpsO* promoter from *M. tuberculosis* (*Mtb*) was created analogously. In this case, the promoter region was 5’-extended up to position −158 from TSS. Primers used for the two-step PCR: Mtb_rpsO-for (5’-AGA*ACGCGT*TCGAATCGGTGCG, MluI site italicized), Mtb_rpsO-egfp-rev (5’-CGCCCTTGCTCACGAAATGTCTCCATC), Mtb_rpsO-egfp-for (5’-GATGGAGACATTTC**GTG**AGCAAGGGCG, initiator GUG in bold) and pQEegfp-rev (see above). All amplification reactions were done with the Tersus Plus PCR kit (Evrogen).

### Creating the plasmid for ectopic expression of the *Msm rpsO* gene in *M. smegmatis*

To create the pAMYC derivative expressing *Msm* S15, we amplified the *rpsO* gene flanked with the 5’-extended promoter and terminator regions on *Msm* genomic DNA by using Q5 DNA polymerase and primers P(−231)rpsO-for bearing BamHI (5’-TGA***GGATCC***AATCCGACGTTCTC, BamHI in bold italicized) and Msm-rpsO-rev (5’-ACT***AAGCTT***GCATGTCCGCAGAC, HindIII in bold italicized). The PCR product was treated with BamHI /HindIII and then ligated into pAMYC treated with the same endonucleases. The ligation mix was used to transform *E. coli*; plasmids were isolated from Cm-resistant colonies, sequenced and further used to transform *M. smegmatis* bearing the reporter *Msm (or Mtb) rpsO-egfp*.

### Cell growth and GFP assay

Transformation-proficient *M. smegmatis* mc^2^ 155 (53) was used for electroporation with pMV306 (Kan^r^) derivatives bearing the *rpsO-gfp* reporter genes to provide their insertion into the chromosome. The Kan^r^-transformants were selected on LB-Kan agar plates, and then used for competent cell preparation and electroporation with an empty shuttle vector pAMYC or with its derivative carrying the *Msm rpsO* gene for S15 expression *in trans* (see above). The transformants were selected on LB-Kan-Cm agar plates, then grown at 37°C in LB supplemented with 34 μg/ml Cm and 0.05% Tween 80 (to prevent cell clumping) and harvested in exponential phase (OD_600_ ∼0.7-0.8). Protein extracts were prepared as described (54) with slight modifications. Cell pellets were resuspended in PBS and broken by using Beat Beater and 0,1 mm zircon beads (BioSpec Products) (3 times for 30 s on ice, with 1:4 vol/vol ratio of beads to cell suspension). The cell lysates were clarified by centrifugation (20 min, 12000 rpm at 4 °C), supernatants were treated with RQ-DNase (Promega) for 30 min on ice and used for the eGFP assay. Protein concentration in clarified lysates was determined by Bradford assay (Bio-Rad). EGFP fluorescence in protein samples was measured in a 96 well microplate using Tecan Genios Pro fluorescence microplate reader (Tecan, Switzerland). The results were normalized to the protein concentration in samples. Usually, each sample was obtained at least in three biological replicates. As a background control, we used protein lysates obtained from *Msm* exponentially grown cells.

## Acknowledgements

We thank Dr. Elena Salina for a gift of genomic DNA from *M. tuberculosis*. We are grateful to Dr. Tatyana Azhikina, Dr. Yulia Skvortzova and Artem Grigorov for their advices and helpful discussion. This work was supported by the RFBR grant №18-04-00743.

